# Principles of studying a cell - a non-boastful paper for all molecular biologists

**DOI:** 10.1101/027680

**Authors:** Han Chen, Xionglei He

## Abstract

Studies of a cell rely on either observational approaches or perturbational/genetic approaches to define the contribution of a gene to specific cellular traits. It is unclear, however, under what circumstances each of the two approaches can be most successful and when they are doomed to fail. By analyzing over 500 complex traits of the yeast *Saccharomyces cerevisiae* we show that the trait relatedness to fitness determines the performance of observational approaches. Specifically, in traits subject to strong natural selection, genes identified using observational approaches are often highly coordinated in expression, such that the gene-trait associations are readily recognizable; in sharp contrast, the lack of such coordination in traits subject to weak selection leads to no detectable activity-trait associations for any individual genes and thus the failure of observational approaches. We further show that genetic approaches can be successful when the genes responsible for coordinating the target genes of observational approaches are perturbed. However, because the system-level cellular responses to a random mutation affect more or less every gene and consequently every trait, most genetic effects convey no trait-specific functional information for understanding the traits, which is particularly true for traits subject to weak selection.

**Significance statement:** Cell research is nearly exclusively based on empirical data obtained through either observational approaches or perturbational/genetic approaches. It is, however, increasingly clear that an analytical framework able to guide the empirical strategies is necessary to drive the field further ahead. This study analyzes ~500 complex traits of the yeast *Saccharomyces cerevisiae* and reveals the organizing principles of a cell. Specifically, a cell can be viewed as a factory, with each trait being the product of a production line operated directly by workers who are supervised by managers. For a cellular trait produced by many workers, the coordination level of the workers determines the performance of observational approaches. Meanwhile, the coordination of workers is realized by managers that are recruited and/or maintained by natural selection. Thus, observational approaches are expected to fail for traits subject to little selection, and genetic approaches can be successful only when the managers of fitness-tightly-coupled traits are perturbed. The manager-worker architecture built by natural selection explains well the origins of global epistasis and ubiquitous genetic effects, two major issues confusing current genetics and molecular and cellular biology, providing a clear guideline on how to study a cell.

## Introduction

Understanding a cell is to characterize the genes underlying each cellular trait. There are currently two basic strategies in cell research: 1) observational approaches that relate a gene to a trait based on statistical associations of the trait with the gene's activity; 2) perturbational (or genetic) approaches that relate a gene to a trait by the effect on the trait after perturbing the gene^1^. Technical advances in recent years enable genomic profiling of various types of gene activity (e.g., mRNA level, protein abundance, protein phosphorylation, protein location, protein-protein interactions, protein-DNA/RNA interactions), greatly facilitating observational approaches to inferring gene-trait associations. Meanwhile, genome-wide reverse genetic screenings based on homologous recombination^2^, RNAi^3^ or CRISPR-Cas9^4^ are designed to reveal the whole set of genes whose perturbations alter a trait. It is thus increasingly clear that data acquisition is no longer a major hurdle to understanding a cell. However, three key challenges remain in the field. First, the performance of observational approaches is heavily compromised by between-gene epistases that appear to be pervasive^5^. Second, because all genes are connected with each other in a cell to influence traits, perturbation of any one gene could, in principle, propagate through the cellular system to affect any trait to some extent^6^. Because no functional insight can be gained from claims of a gene responsible for all traits or a trait affected by all genes, the rationale for using genetic approaches to understand specific traits is unclear. Third, the gene-trait associations revealed by observational approaches are often not replicated using genetic approaches and vice versa.

To address the three issues in this study we analyzed 501 morphological traits that are measured in 4,718 single-gene deletion mutants of the yeast *Saccharomyces cerevisiae*^7^. We identified a single dominant factor that determines the performance of observational approaches in revealing gene-trait associations. We then compared the functional properties of the genes revealed by observational approaches and genetic approaches, and developed a model to explain why the gene-trait associations identified by observational approaches have little overlap with those identified by genetic approaches. In the end, we showed that only in limited circumstances can the genetic effects on a trait provide specific functional information for understanding the trait.

## Results

### The performance of observational approaches varies dramatically in different traits

We used mRNA level as the representative gene activity to test the performance of observational approaches in revealing the genes associated with a trait, by taking advantage of up to ~1,500 microarray-based expression profiles of the yeast single-gene deletion mutants^8^ (Fig. S1). We identified for each trait the *<u>e</u>*xpression *<u>i</u>*nformative *<u>g</u>*enes (EIGs) whose expressions are linearly correlated to the trait in a robust fashion (Methods). The number of EIGs found for a trait varied substantially, ranging from zero to ~1,000. Interestingly, traits with fewer EIGs did not necessarily show a simpler genetic architecture, which is measured by the number of *<u>g</u>*enetically *<u>i</u>*nformative *<u>g</u>*enes (GIGs) that, when deleted, show statistically significant effects on the traits^9^; for example, there were on average 129±22.9 GIGs for traits with <10 EIGs and 167±15.6 for the rest traits with ≥ 10 EIGs (*p* > 0.05, Mann-Whitney U test; Fig. S2). Given that a trait often has >100 GIGs, there should be a large number of genes whose altered expression causally mediates the diverse genetic effects on the trait. Because some of the identified EIGs may mediate the genetic effects while the others may be reactive to their traits^10,11^, we examined the expression-trait correlations of 118 EIGs that were also found in ~60 F1 segregants of a hybrid of two *S. cerevisiae* strains^12^. After analyzing the segregation patterns of traits, QTLs and gene expression in the F1 segregants (Methods), we estimated that 15-40% of the EIGs can causally affect their traits (Fig. S3).

### The performance of observational approaches is determined by the coordination of the target genes

In principle, with an increasing number of causal factors the variance explained by each factor would become negligible. It is thus surprising to observe in a single trait hundreds of EIGs that each show a significant expression-trait correlation. A reasonable explanation is that the expression regulation of these EIGs is highly coordinated such that they form a small number of independent controllers. The lack of EIGs in the remaining traits cannot be explained by a small number of causal genes, because the variance explained by any individual EIG in traits with few EIGs was minimal (Fig. S4). Therefore, there must be many causal genes to mediate the diverse genetic effects on each of the traits. Observational approaches attempt to uncover these causal genes, but fail to do so because they function in an uncoordinated fashion, resulting in pervasive gene-gene interactions (or epistasis), including antagonistic epistasis ^13^, and thus no detectable expression-trait associations for individual genes. To illustrate this reasoning, we simulated a scenario in which a trait is affected by 50 genes in an additive fashion (Methods). As the co-expression of the 50 genes decreases, the probability that an individual gene remains significant expression-trait correlation diminishes quickly (Fig. S5A). The same effect size corresponding to both up- and down-regulation of a focal gene, a phenomenon often explained by invoking antagonistic epistasis^13^, became common when the co-expression was minimal (Fig. S5B). Note that this pattern is not merely the product of our specific simulation; rather, it is expected given that the variance explained by each individual factor will be small when the number of independent causal factors is large.

### Natural selection underlies the coordination of the target genes of observational approaches

We reasoned that such coordination must be built and/or maintained by natural selection and thus predict the failure of observational approaches in traits subject to little selection. We used cell growth rate as a proxy of fitness in single-celled yeast and calculated the trait relatedness to fitness for each of the morphological traits (Methods). Remarkably, the number of EIGs found in a trait was largely explained by its trait relatedness to fitness (Spearman’s ρ = 0.89, n = 501, *p* < 10^−16^). There were typically several hundred EIGs in a trait tightly coupled with fitness but no EIGs at all in those with no significant correlation to fitness (Fig. 1). The two orders of magnitude difference in the total EIG number suggested that the disparity between the fitness-coupled and fitness-uncoupled traits is robust despite the contamination of reactive EIGs (Fig. S3). This pattern was also observed when 57 largely unrelated exemplar traits with divergent EIG compositions were considered (Methods) (Fig. S6). Further the pattern cannot be explained by noise in trait measurement (Fig. S7) or by a smaller variation in fitness-uncoupled traits (Fig. S8). Thus, natural selection is required for building and/or maintaining coordination of the target genes of observation approaches, resulting in robust gene expression-trait correlations observed in fitness-coupled traits. In sharp contrast, a lack of selection constraints on fitness-uncoupled traits results in poor coordination of the target genes, leading to global epistases that strictly prevent observational approaches from revealing gene-trait associations. Note that for simplicity throughout the manuscript fitness-coupled (-uncoupled) traits refer to those whose trait value is tightly (loosely) coupled with cell growth rate; we are fully aware that, strictly speaking, all traits are fitness-coupled to some extent.

**Fig. 1.**
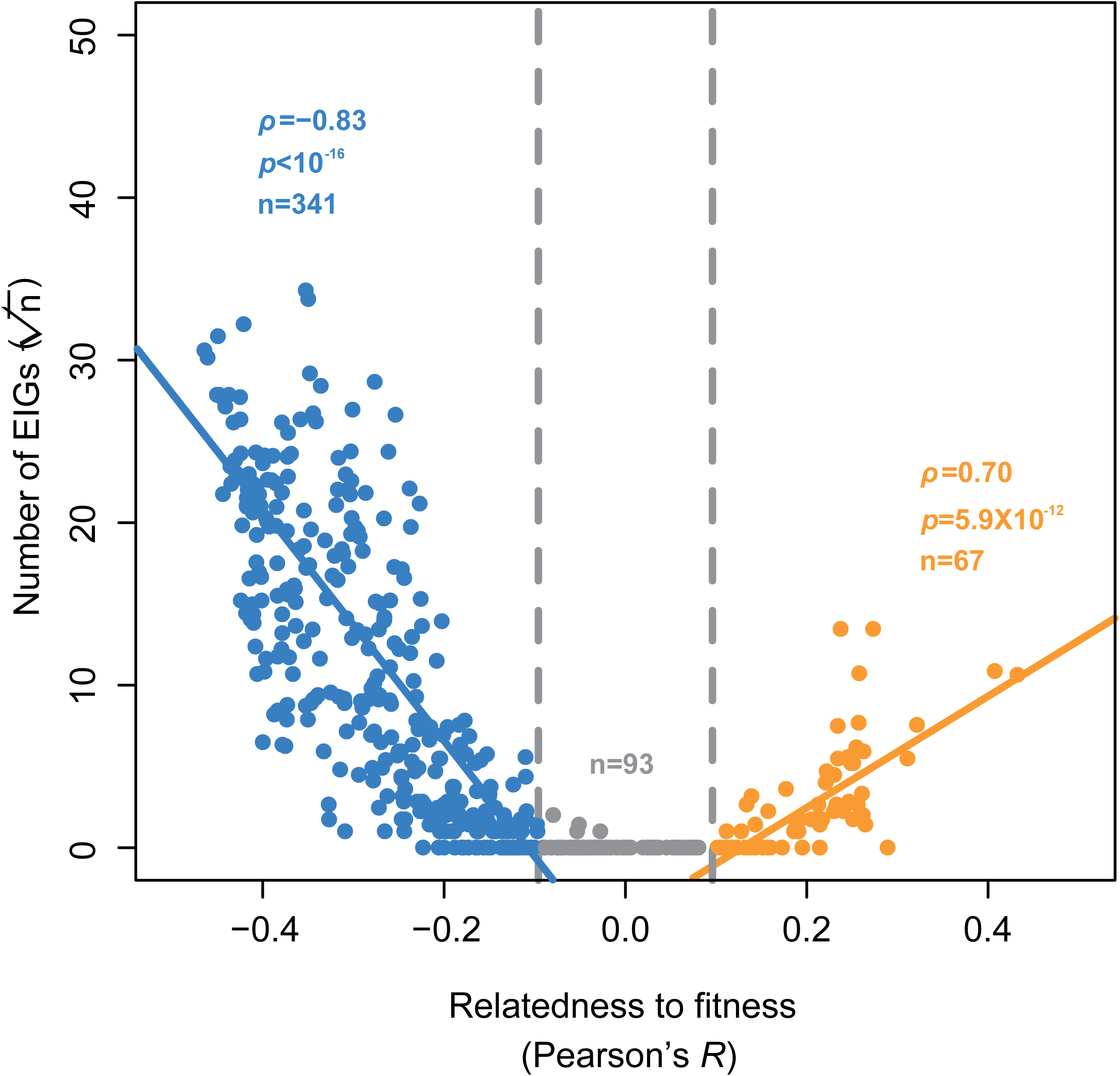
The number of EIGs found in a trait is dependent on trait relatedness to fitness. The y-axis shows the square root of the number of EIGs, and the x-axis is the trait relatedness to fitness measured by the Pearson’s *R* between the trait values and the cell growth rates of the yeast mutants, with *R* > 0.1 or *R* < −0.1 regarded as statistically significant after controlling for multiple testing. Each dot represents a trait, and ρ shows the Spearman’s correlation coefficient.

### Understanding a super-complex trait using EIG-modules

How well can we understand a trait using its EIGs? We tested this issue by examining cell growth rate, the yeast fitness-determining trait with arguably the most complex genetic architecture, as evidenced by the fact that over one third (~2,000) of the yeast genes, when deleted, show a growth rate reduction greater than 5% in the rich medium YPD^14^. Using the functional data considered above, we identified over 900 growth rate related EIGs using a stringent criterion; these form six protein modules that each have a clear Gene Ontology enrichment (Table S1) (Methods). Analysis of the six EIG-modules revealed a variety of novel mechanistic insights into the regulation of yeast cell growth (Supplementary Note 1 and Fig. S9). Also, a simple linear function integrating the six EIG-modules explained up to ~50% of the growth rate variation of over 400 mutants (Pearson’s *R* = 0.69, n =442, *p* < 10^−16^; Fig. 2A). Note that the cell growth rates considered here are measured using the Bar-seq technique^14^, which is believed more accurate than the microarray-based method^15^ or colony-size-based method^5^, both used previously for quantifying growth rates of the yeast mutants. Using the same set of mutants, we showed that the Pearson’s *R* is 0.77 between the microarray-based measures and the Bar-seq-based measures, and 0.63 between the colony-size-based measures and the Bar-seq-based measures (Fig. 2B and C), suggesting that the EIG-module-based linear model was comparable to the two conventional experimental approaches in estimating yeast cell growth rate.

**Fig. 2.**
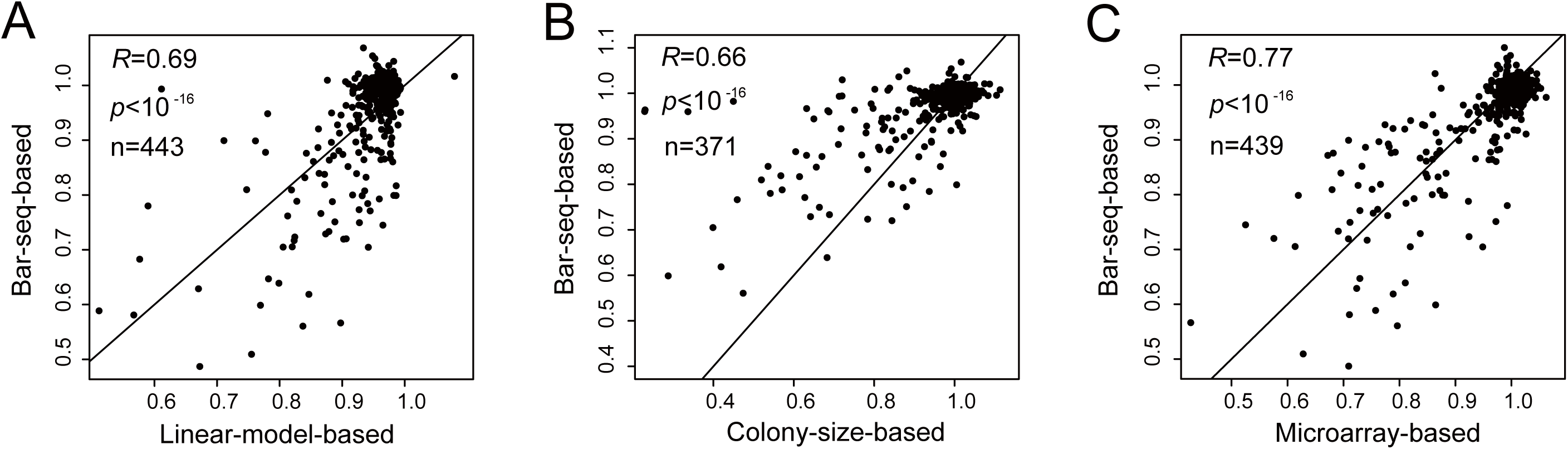
Good performance of EIGs in modelling a super-complex trait. **(A)** Growth rates of the yeast mutants based on the Bar-seq technique or the linear model written as *G* = -1.740ED*_M1_* - 0.435ED*_M2_* - 0.725ED*_M3_* - 0.071ED*_M4_* + 0.794ED*_M5_* - 0.058ED*_M6_* + 1.019, where *G* stands for growth rate. Each dot represents a deletion mutant, with the Pearson’s *R* shown. **(B)** Growth rates of the yeast mutants measured by the Bar-seq technique or the colony-size-based method, with 72 mutants excluded due to the lack of the colony-sized-based measures. **(C)** Growth rates of the yeast mutants measured by the Bar-seq technique or the microarray-based method, with four mutants excluded due to the lack of the microarray-based measures.

### A manager-worker model explaining the disparity between observational and genetic approaches

In this study, EIGs are revealed by observational approaches and GIGs by genetic approaches. The number of EIGs does not predict the number of GIGs (Fig. S2); also, there are no more overlaps than expected by chance (*q* = 0.1) between EIGs and GIGs of the same traits in the 109 traits that each have ≥ 10 EIGs and ≥ 10 GIGs (Fig. S10). A close examination showed that, compared with EIGs, GIGs tend to be those that, when deleted, affect a large number of genes’ expressions but themselves are less responsive to genetic perturbations (Fig. 3). It is thus likely that a typical complex trait is responsive directly to the collective activities of a large number of EIGs such that the effects of removing a single EIG are often too small to be detected; removing a GIG affects many EIGs, resulting in generally larger genetic effects that are more detectable. An analogy to this is a production line run by workers and managers. A major productivity slow-down is often due to removing a manager instead of removing a worker, despite the fact that the workers are more directly involved in production. This analysis also helps resolve an important puzzle that genes with expression response to a given condition are often not genetically required for the condition^2,16^.

**Fig. 3.**
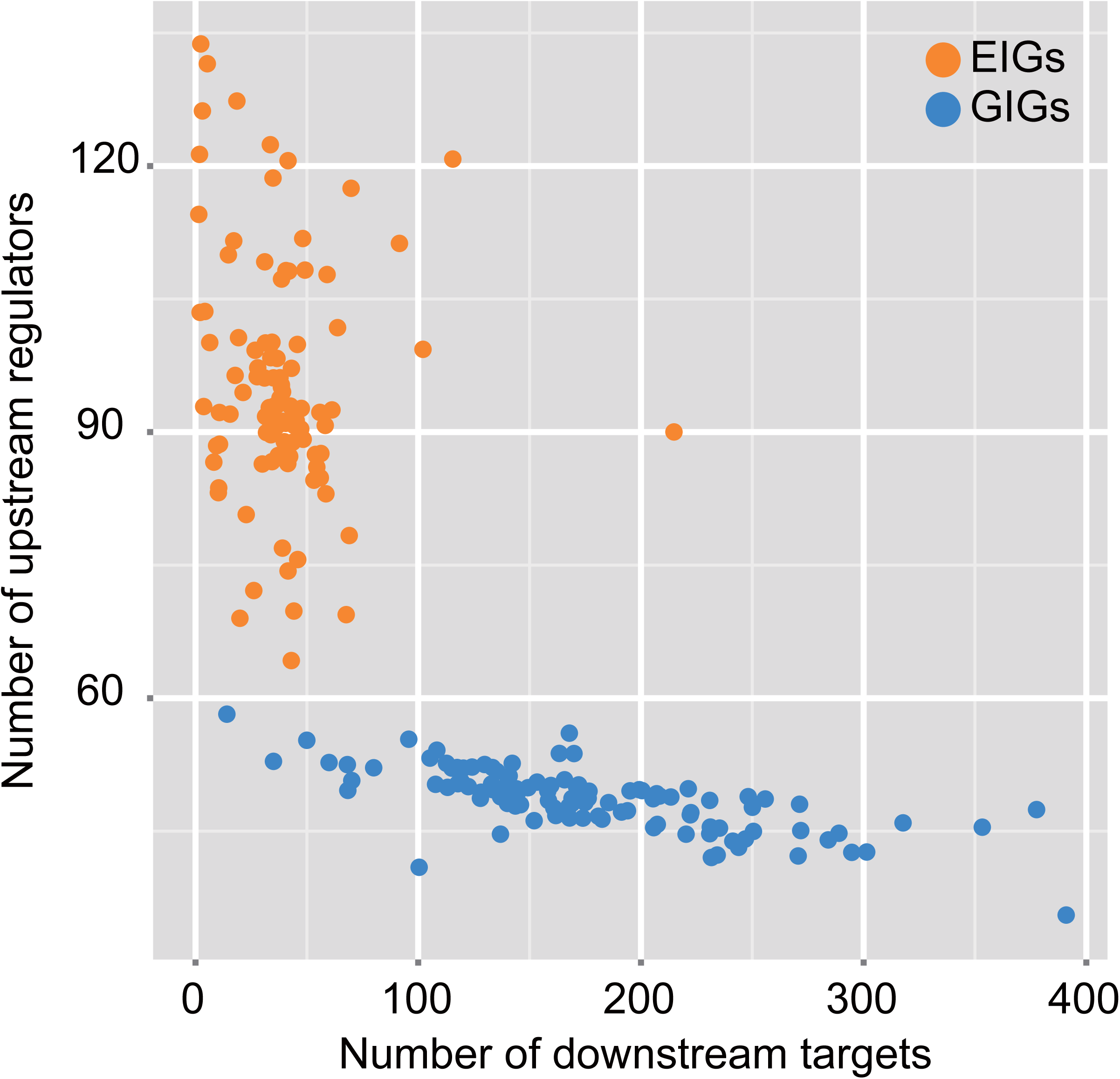
Distinct functional properties between EIGs and GIGs. The numbers of downstream targets (x-axis) and upstream regulators (y-axis) per EIG or GIG. Each dot represents a trait, and the average of all of its EIGs or GIGs is shown for each trait.

### The manager-worker model suggests two types of genetic effects

The other basic strategy in cell research, the genetic approach, is confounded by the potential system-level cellular responses to a mutation, which may alter all traits to some extent. We can use the manager-worker analogy to model the origins of genetic effects. Null mutations on a manager of a trait launch coordinated perturbations onto the related workers, eliciting a trait-specific profile of cellular responses and consequently “*<u>s</u>*pecific” *<u>g</u>*enetic *<u>e</u>*ffects (SGEs). However, system-level cellular responses to a random mutation affect more or less every gene, including the workers but in an uncoordinated fashion, to elicit the non-specific “*<u>u</u>*biquitous” *<u>g</u>*enetic *<u>e</u>*ffects (UGEs). This reasoning suggests two expected differences between SGEs and UGEs. First, SGEs should be found primarily in traits subject to strong selection because recruiting and/or maintaining managers to coordinate workers requires natural selection. SGEs might also be generally stronger than UGEs because of the coordinated changes of workers. Second, SGEs can be used to identify workers of the focal trait because of the profile of coordinated changes that exposes the workers, while UGEs will not exhibit such coordinated changes and thus provide no trait-specific functional information.

### A small number of disproportionately large genetic effects found in fitness-coupled traits

We modeled for each trait the per-gene effect size with the commonly-used Gaussian function that is expected to capture the size distribution of UGEs^17^, which is continuous due to the system-level cellular responses to random mutations (Methods). We used quantile-quantile plot to compare the Gaussian approximation to the true distribution and found that the two distributions often fit each other reasonably well (Fig. 4A and B). In some traits, however, there were disproportionately large effects that are far beyond the Gaussian approximation (Fig. 4C and D). We thus defined outlier effects as those with absolute Z-scores > 5.06, which corresponds to *p* = 2.12 × 10^−7^ in the standard Gaussian distribution or *q* = 0.001 after the Bonferroni correction for multiple testing (*q* = *p* × 4,718). The number of outliers identified in a trait varied from zero to ~50. Interestingly, the outlier number of a trait was highly correlated with the trait relatedness to fitness, and there were often a negligible number of outliers in traits with no significant correlation to fitness (Fig. 4E). This pattern remained when the Z-score cutoff of defining outliers was changed to 4.56 (*q* < 0.005) or to 4.06 (*q* < 0.01) (Fig. S11), or when only uncorrelated exemplar traits were analyzed (Fig. S12). Because outlier effects may cause strong fitness coupling of a trait, we recalculated for each trait its relatedness to fitness after excluding the outlier genes (Methods). The recalculated values were highly correlated to the original ones (Pearson’ *R* = 0.96, n = 501, *p* < 10^−16^; Fig. S13), suggesting that it is fitness coupling that determines the presence of outliers. According to our reasoning above, it is likely that the outliers represent SGEs and the non-outliers are UGEs.

**Fig. 4.**
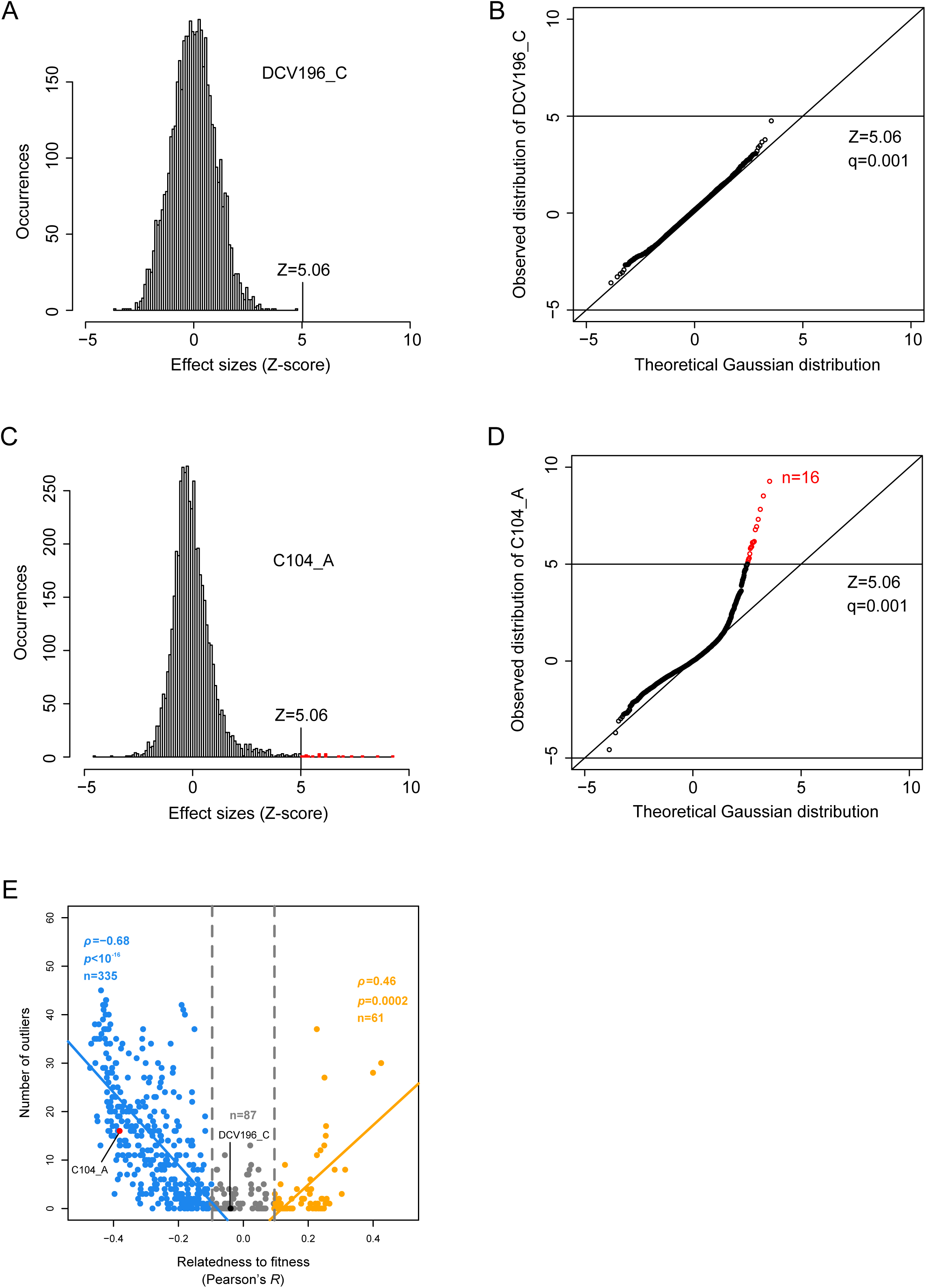
Natural selection determines the outlier genetic effects. The frequency distribution of effect sizes **(A)** and the Q-Q plot comparing this distribution with its Gaussian approximation **(B)** in the trait DCV196_C. The frequency distribution of effect sizes **(C)** and the Q-Q plot comparing this distribution with its Gaussian approximation **(D)** in the trait C104_A. **(E)** The number of outliers found in a trait is highly correlated to the trait relatedness to fitness that is measured by Pearson’s *R* between trait value and cell growth rate of the yeast mutants, with *R* > 0.1 or *R* < -0.1 regarded as statistically significant after controlling for multiple testing. Each dot represents a trait, and p shows the Spearman’s correlation coefficient.

### Trait-specific functional information provided by SGEs but not UGEs

We expect SGEs (but not UGEs) to convey trait-specific functional information. This hypothesis can be tested using gene expression profiles of the yeast mutants. To avoid potential false positives we focused on the GIGs whose deletion effects are statistically significant under a stringent cutoff^9^. There were typically a few hundred GIGs found in a trait no matter whether the trait is highly related to fitness or not (Fig. S14). We examined in each trait the top 20 GIGs with the largest effects that also have available expression profiles. This included typically ~10-18 outlier GIGs, which correspond to SGEs, in fitness-coupled traits but only non-outlier GIGs, which correspond to UGEs, in fitness-uncoupled traits (Fig. 5A). We calculated for each trait the expression profile similarity between the top 20 GIG mutants (Methods). The resulting expression similarity for a typical fitness-uncoupled trait was not stronger than the background (Fig. 5B), which was measured by comparing all GIGs of the 129 different traits analyzed here (Methods). As predicted, even the strongest UGEs show no trait-specific cellular signature.

**Fig. 5.**
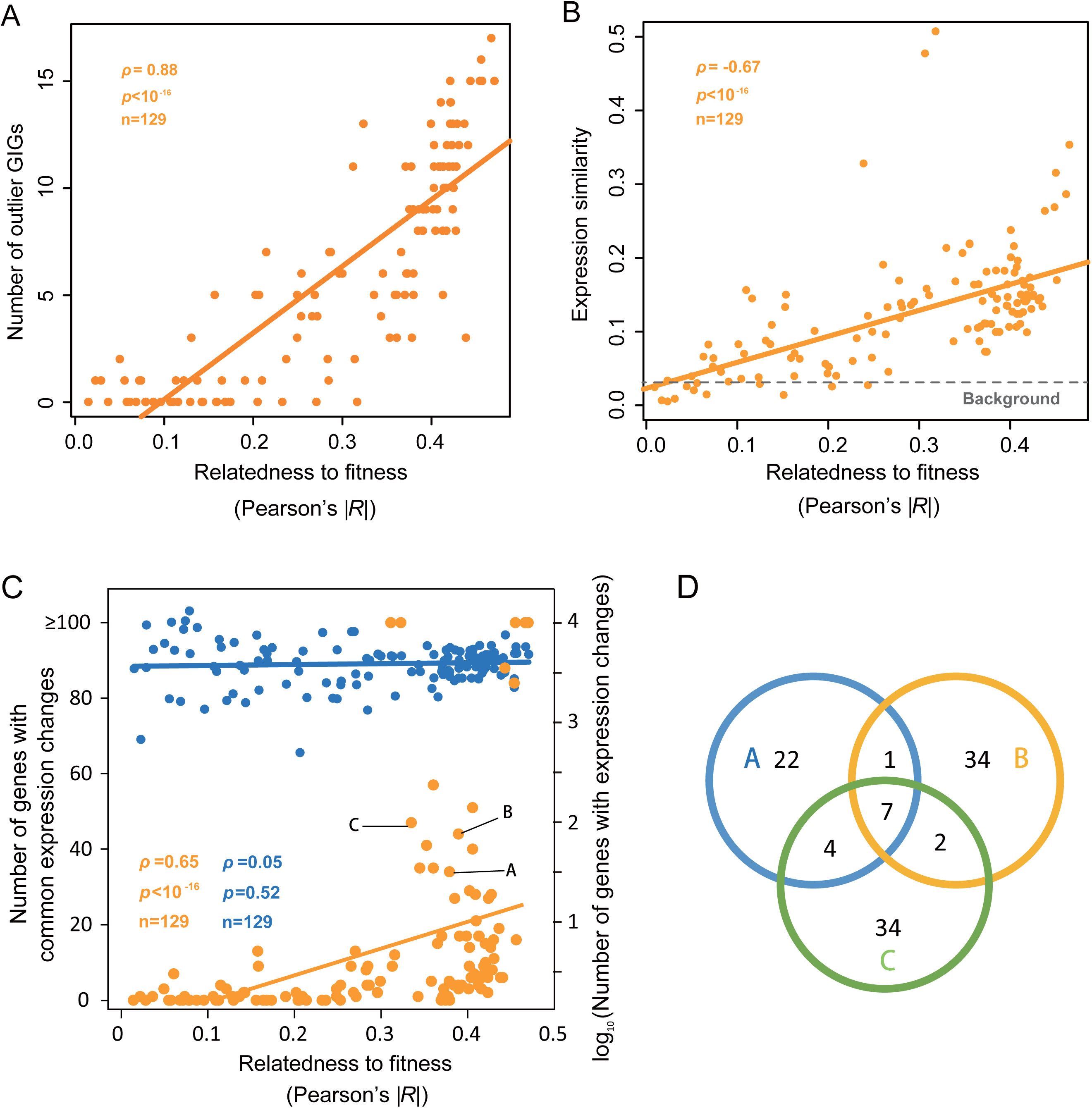
Characterization of SGEs and UGEs. **(A)** The number of outliers among the top 20 GIGs of a trait as a function of the trait relatedness to fitness. Each dot represents a trait. **(B)** The expression similarity among the top 20 GIG mutants of a trait as a function of the trait relatedness to fitness. Each dot represents a trait, and expression similarity is the average Pearson’s *R* of all 190 mutant pairs. The background is the average expression similarity between all top 20 GIG mutants of the 129 traits. **(C)** The number of genes with common expression changes (yellow) and the total number of expression changes (blue) in mutants of the top 20 GIGs as a function of the trait relatedness to fitness. Traits C104_C, D117_C and D134_C are labeled A, B, and C, respectively. **(D)** Plenty of trait-specific commonly responsive genes in the three representative traits highlighted in panel C.

We observed much stronger between-mutant expression similarity for the fitness-coupled traits (Fig. 5B). This pattern could be due to either fewer total expression changes or more shared expression changes. We identified the genes that are commonly down- or up-regulated in the top 20 GIG mutants of each trait under a statistical cutoff where the expected number of such genes is slightly smaller than one (Methods). Despite the fact that total expression changes were similar between fitness-coupled and -uncoupled traits, fitness-coupled traits typically had a few dozen genes with common expression changes but most fitness-uncoupled traits had no such genes (Fig. 5C). The absence of such commonly responsive genes in the fitness-uncoupled traits further supported the notion that no trait-specific cellular responses cause the UGEs. Among the 1,060 non-redundant commonly responsive genes identified in the 129 traits, the mean and median number of traits a gene involved are 2.97 and 2, respectively, suggesting plenty of trait-specific functional information provided. For example, a close examination of three representative fitness-coupled traits revealed a distinct composition of commonly responsive genes in each trait (Fig. 5D). Interestingly, approximately 60% (8.7-fold enrichment with the 95% confidence interval of 5.6~12.2-fold; permutation test) of the commonly responsive genes are also expression informative genes (EIGs) of the same traits. Notably, 15-40% of EIGs are the workers whose activities directly determine the traits (Fig. S3). These data suggested that the outlier GIGs are the managers recruited and/or maintained by natural selection to coordinate the related workers. The expression responses to perturbing a manager of a fitness-coupled trait expose the workers of the trait, justifying the use of genetic approaches to studying traits of this type.

## Discussion

There are three caveats that warrant discussion. First, among the many types of gene activities, only mRNA level was examined because of data availability. Although there are differences between mRNA level and protein activities, the general conclusions, for example, that the performance of observational approaches is dependent on the coordination of workers, do not seem to be sensitive to the gene activity considered. Second, the cell growth rate measured in YPD is not ideal for representing the natural fitness of yeast, although the relative growth rates of the deletion mutants measured in diverse media are largely correlated^15^. This potential problem, however, is unlikely to generate the striking differences observed between the fitness-coupled and -uncoupled traits; it would instead blur the comparison to make our findings more conservative. Third, because the genetic/phenotypic space represented by the F1 segregants of the BY x RM hybrid is limited, only 118 identified EIGs were tested for their causal effects on the traits, which gave a rough estimation of the proportion of causal EIGs (i.e., workers of a trait). Sampling more variations in natural populations would give a more accurate estimation, but a refined estimate is unlikely to overturn our conclusion that a significant proportion of EIGs are causal.

This study reveals the organizing principles of a cell: A cell can be viewed as a factory, with each trait being the product of a production line operated directly by workers who are supervised by managers. For a complex trait produced by many workers, the coordination level of the workers determines the performance of observational approaches; specifically, the associations between individual workers and the trait are readily recognizable when the workers’ activities are coordinately changed. Meanwhile, the coordination of workers is realized by managers that are recruited and/or maintained by natural selection, so genetic approaches can be successful only when the managers of a fitness-coupled trait are perturbed, which generates a trait-specific profile of cellular responses to expose the workers.

Complexity arises from the absence of such coordination. Although current genetics is defined by statistics (Supplementary Note 2 and Fig. S15), the fact that all genes are connected in the cellular network predicts that perturbing a random gene would affect all or nearly all genes including the workers and consequently the focal traits. Such ubiquitous genetic effects provide little information necessary for revealing the workers because the workers are unlikely changed coordinately and thus generate trait-specific profiles, predicting the failure of genetic approaches in this context. The lack of coordination among workers suggests complex between-worker epistasis underlying traits, predicting the failure of observational approaches. Thus, most, if not all, confusions in current genetics and molecular and cellular biology can be ascribed to the ubiquitous genetic effects and global epistases, both of which result from the lack of effective selection. This notion is particularly important for human biology, because natural selection is inefficient in humans due to the small effective population size^18^, and because aging-associated diseases or traits are often of little fitness relevance but of high interest to researchers^19,20^. One may argue that such global epistases and ubiquitous genetic effects are exactly the challenges we need to address, but the lack of selection constraint predicts that they might be *ad hoc* phenomena sensitive to genetic and environmental backgrounds^21^. A robust discussion of both the strategy and the necessity of studying these issues is needed.

## Methods

### Data

The yeast *Saccharomyces cerevisiae* single-gene deletion stock was generated by Giaever et al. (2002), with 4,718 mutant strains each lacking a nonessential gene being considered in this study. As for cell growth rates of the above mutants measured in the rich medium YPD (yeast extract, peptone, and dextrose), the Bar-seq-based data were by Qian et al. (2012), the microarray-based by Steinmetz et al. (2002), and the colony-size-based by Costanzo et al. (2010). The 501 morphological traits of the mutants (SCMD) were characterized by Ohya et al. (2005), and the genetically informative genes (GIGs) that show significant phenotypic effects after deletion were defined for 220 traits by Ho and Zhang (2014), with 216 reproducible using the updated data in SCMD and thus included in this study. The microarray-based expression profiles of 1,484 deletion mutants were generated by Kemmeren et al. (2014); gene A was called the downstream target of gene B and B the upstream regulator of A, if gene A shows a significant expression change (*P* < 0.0001 as provided in the original data) in the gene B deletion mutant.

### Identification of expression informative genes (EIGs)

Nearly all of the 501 morphological traits of the deletion mutants show a bell-shape distribution, with the median trait value very close to that of the wild-type (Fig. S16). There are 1,328 strains with both the expression profiles generated by Kemmeren et al. (2014) and the Bar-seq-based cell growth rates. We randomly divided the 1,328 yeast strains into two sets, with two thirds (885) for Set #1 and one third (443) for Set #2. There are 6,123 yeast genes on the chip used by Kemmeren et al. (2014). We first generated 500 artificial datasets, each containing 443 strains picked randomly from the 885 Set #1 strains with replacements. We calculated the Pearson’s *R* between expression level and trait value for each of the 501 x 6123 trait-gene pairs in each the 500 artificial datasets, respectively. The *R* values were then transformed into *p*-values using T-test; in each trait under examination we thus obtained 500 *p*-values for each gene. We defined the correlation robustness (*r*-value) of a given gene as the harmonic mean of its *p*-values after dropping both the highest and the lowest 5% of its 500 *p*-values, which was then multiplied by 6,123 for multiple testing correction. Genes with the corrected *r*-values < 0.01 were considered as potential expression informative genes (EIGs). To further reduce false positives, we required that the potential EIGs also show significant expression-trait correlation in the independent Set #2 mutants, resulting in a total of 2,541 non-redundant genes identified as EIGs of at least one trait, with and mean and median number of traits an EIG affects being 27 and 11, respectively.

Morphological traits are not independent; for instance, the size and the diameter of a cell are correlated. To reduce correlated traits, we employed an un-supervised affinity propagation strategy proposed by Frey and Dueck (2007) to cluster the 501 traits based on the *r*-values of all genes, resulting in 57 clusters each with an exemplar trait.

The distribution of cell growth rates of the mutants is highly biased, with the majority close to the rate of the wild-type. We thus computed the expression-growth rate correlation using the univariate Cox’s regression model that emphasizes the difference of two categories, with growth rate as the parameter “time”, strains of growth rate <0.9 weighted as “event = 1”, and all others as “event = 0”. Specifically, we performed the Cox’s regression analysis using the 500 artificial datasets described above and obtained 500 *p*-values for every yeast gene. The corrected *r*-value was computed as previously described and a total of 911 genes each with the corrected *r*-value < 0.001 were defined as expression informative genes (EIGs) of the cell growth rate. We found that the Cox’s regression is more conservative than the Pearson’s regression in defining EIGs. The 911 EIGs identified in the Set #1 mutants were assembled into protein modules and tested for their performance in modelling the cell growth rate using the independent Set #2 mutants.

### Determination of causal associations between EIG expression and traits

Information of the genotype, expression and morphology of 62 F1 segregants of a hybrid of two yeast strains (BY4716, a derivative of S288c, and YEF1946, a derivative of RM11-1a) was obtained from Nogami et al. (2007), with three segregants excluded from further analyses because of unmatched IDs. Because there is no major difference between the two parental yeast strains in most of the morphological traits, there are only 118 EIGs whose expression-trait correlations were also detected in the 59 F1 segregants with *q* < 0.01 (two-tailed T test with Bonferroni correction for multiple testing). The causality of the EIG-expression versus trait association was resolved using the Network Edge Orienting (NEO) method developed by Aten et al. (2008). Following the manual provided by NEO, we calculated the LEO.NB.CPA score and the LEO.NB.OCA score with all genotype information (SNPs) inputted; for each association the two causality directions (i.e., EIG-expression -> trait and trait -> EIG-expression) were tested separately. We defined a cause association if the LEO.NB.CPA score > 0.8 and the LEO.NB.OCA score > 0.3, which corresponds to a false discovery rate of 0.05. We found 18 EIG-expression -> trait and 27 trait -> EIG-expression causal associations, but failed to assign a reliable causal association for the rest 118-18-27=73 associations. Thus, the proportion of causal EIGs is 18/(18+27) = 40% (or 18/118 ~= 15% by assuming no positive in the 73 uncertain associations).

### Modelling the effects of reducing the coordination of causal factors

Suppose there is a trait controlled by 50 genes, and the expression level of each gene relative to the wild-type follows the standard normal distribution. The trait value is defined as the average relative expression level of the 50 genes plus a random number drawn from the standard normal distribution. For a given co-expression (or coordination) level of, say, 0.5, we simulated 1,000 expression profiles where the average Pearson’s *R* of all gene pairs is 0.5, and the resulting trait values typically follow an approximately normal distribution with mean equal to zero, the wild-type trait value. To what extent the relative expression level of an individual causal gene can predict the trait value defined by all the 50 genes (plus error) is then examined.

### Calculation of the relatedness of the morphological traits to fitness

The relative cell growth rate is a reasonable measure of the relative fitness for the single-celled yeast. Because in this study all cellular traits are measured in YPD, we used the cell growth rate in YPD as the proxy of fitness.

Given the bell-shape distribution of a morphological trait where the wild-type trait value is almost always located in the middle, both increase and decrease of a trait value relative to the wild-type could affect fitness in the same direction. Thus, we divided for a given trait the 4,718 mutants into two equal halves according to the trait values, and calculated the Pearson’s *R* between trait value and fitness for each half of the mutants separately, resulting in two *R*s for every trait. The *R* with the larger absolute value was used to represent the relatedness of the trait to fitness. To assess the effects of outliers on the estimation of fitness coupling, we removed for each trait the top 50 trait values from each side and recalculated the trait relatedness to fitness. We also computed the Pearson’s *R* without separation of the mutants into two halves, and found that it is often highly similar to the relatedness obtained above (Fig. S17).

### Assessment of effects of trait measurement

To characterize the yeast morphological traits Ohya et al. examined on average 400 individual cells for each mutant. The trait value of a given mutant is the mean trait value of the examined cells. Despite the generally large number of examined cells, for some traits there were only a few tens of informative cells, which may affect the reliability of the measurements. To address this issue, we randomly divided the examined cells of each mutant into two equal halves and computed the traits for each half separately. For each trait we then computed the Pearson’s *R* between values derived from the first half and values from the second half. The consistency between the two halves varies substantially among the traits, but is not dependent on the trait relatedness to fitness.

### Calculation of expression distance (ED)

For a given EIG module its expression distance (ED) between a mutant and the wild-type was defined as the normalized Euclidian distance between the two expression profiles:

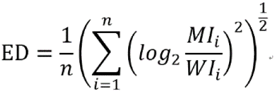
where *MI*_i_ and *WI*_i_ are the expression level of the i^th^ gene in the mutant and wild-type strains, respectively, and n is the number of genes in the module.

### Separation and functional annotation of EIG modules

Yeast protein-protein interactions (PPIs) were downloaded from BioGrid, a database built by Stark et al. (2006). For a given trait we constructed a non-directional, unweighted PPI network composed exclusively of its EIGs. Protein modules were separated using an order statistics local optimization method (OSLOM) proposed by Lancichinetti et al. (2011) with default settings. To annotate the biological functions of these protein modules, we performed the gene ontology (GO) enrichment analysis for each module using BinGO by Maere et al. (2005) and Cytoscape by Shannon et al. (2003). We obtained seven modules formed by the EIGs associated with cell growth rate, among which six were found to be enriched with functionally similar proteins under a false discovery rate of 0.001 (Table S1).

### Characterization of two types of genetic effects

We first examined the distribution of raw trait values for each of the 501 morphological traits, and excluded 18 traits whose distribution is not uni-modal (*p* < 0.05, Hartigan’s Dip-test), leaving 483 traits for further analyses. We normalized the raw trait value *X_ij_* of mutant *j* in trait *i* to Z-score effect size using:

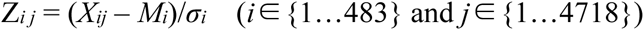
where *M_i_* and *σ_i_* are the mean and standard deviation of the 4,718 mutants in raw trait values of the trait *i*. The outlier effects were defined as the absolute Z-score > 5.06, which corresponds to *p* < 0.001 x 1/4,718 or *q* < 0.001 according to the standard Gaussian distribution.

We defined GIGs with absolute Z-score > 5.06 as outlier GIGs and all others as non-outlier GIGs. For simplicity we excluded 20 traits with a couple of outlier GIGs in the fitness-less-coupled side. There are 1,325 mutant expression profiles available for the expression similarity estimation, so ~27% of GIGs can be studied. For each trait we identified its top 20 GIGs with both the largest effect sizes and available expression profiles in the fitness-coupled side. Sixty-seven traits each with <20 such GIGs were excluded, leaving 196-67=129 traits for further analyses.

Expression similarity between mutants is the average Pearson’s *R* of all pairs of expression profiles of the top 20 GIG mutants of a trait. We compared all top 20 GIGs of the 129 traits to estimate the background between-mutant expression similarity.

For each trait the genes with overall expression up- or down-regulation in the top 20 GIG mutants compared to the other 1,305 mutants were identified as commonly responsive genes of the trait, under the statistical cutoff of *p* < 0.0001 (t-test). Traits C104_C, D117_C and D134_C each with 34, 44, and 47 commonly responsive genes were selected as representative fitness-coupled traits marked with A, B, and C, respectively, in Fig. 5C.

### Estimation of the statistically significant and insignificant effects

There are ~400 individual cells for each mutant and a pool of ~16,000 wild-type cells examined by Ohya et al. (2005), and the trait information of individual cells is available for 216 traits. Because the trait value of wild-type is slightly different from the mean trait value of the 4,718 mutants for most of the traits, all Z-score effect sizes of a trait were adjusted by adding (or subtracting) to ensure that the trait value of the wild-type corresponds to Z = 0. For a mutant with a given adjusted effect size *Z* in a trait, we compared the raw trait values between its 50 randomly-selected cells and 50 random wild-type cells, and used *p*-value < 0.001 (Mann-Whitney U test) to define statistically significant effect. This comparison was conducted for all 4,718 x 216 mutant-trait combinations, and the proportion of significant effects was calculated for all adjusted *Zs* within a given Z-score interval. To estimate the expected proportion of significant effects when the effect size is Z, for a given trait we compared the raw trait values between 50 random wild-type cells and another 50 random wild-type cells each being added (or subtracted) an effect size of *Zσ_i_* (i.e., pseudo-mutants), where *σ_i_* is the standard deviation of the trait for the 4,718 mutants. The same statistical cutoff was applied to define the significant effects, and the proportion of significant effect was derived from 216 traits. This simulation was repeated 100 times to get the confidence intervals. Because variance was difficult to model for the pseudo-mutants with a given mean effect size, we assumed the same variance between the pseudo-mutants and the wild-type population, which would cause strong bias when the given effect size is large. Thus, we limited our analysis to effect sizes ranging from Z = 0 to Z = 0.77, which covers 50% of the data with Z > 0 in a standard Gaussian distribution.

## Acknowledgments

We are grateful to Drs W. Qian, J. Zhang, M. Bakewell, A. Tony, P. Shi and C-I Wu for comments. This work is supported by two research grants from the National Natural Science Foundation of China (#91431103 and #31225014 to X. H.). X.H. is supported also by the Changjiang Scholars Program and the Qin-Nian-Ba-Jian Program. X.H. and H.C. designed the study and wrote the paper; H.C. and X.H. analyzed data.

## Supporting Information of “Principles of studying a cell”

The SI file contains:

Supplementary Note 1 and 2

Table S1

Legends of supplementary figures 1–17

Figures S1-S17

## Supplementary Note 1

Because EIGs presumably function coordinately, we used protein-protein interactions to assemble the ~900 EIGs and obtained six protein modules each with a clear Gene Ontology enrichment (Methods). Interestingly, the six EIG-modules (module-1 to module-6 or M1 to M6) are all related to critical biogenesis processes (Table S1). We computed for each EIG-module its expression distance (ED) between the wild-type yeast and a given mutant (Methods), and examined 87 mutants each with a growth rate less than 80% of the wild-type. With only a few exceptions, these slow-growth mutants formed five clusters (Methods), each corresponding to the expression alterations of distinct modules (Fig. S9A), suggesting that the six EIG-modules represent rather independent causal factors of growth defect, which helped clarify a previous confusion with respect to the distinct effects of ribosome-related genes (M5) and amino acid biosynthesis genes (M2) on the cell growth rates of lab strains and wild strains(*1*). Note that we failed to observe such slow-growth mutant clusters based on the expressions of all individual genes of these modules (Fig. S9B). We conducted partial correlation analysis to reveal potential between-module interaction. Interestingly, the Pearson’s *R* between ED_*M5*_ and the growth rate changed from −0.4 to 0.3 after controlling for the influences of the other modules (Fig. S9C). Because M5 represents ribosomal biogenesis, a process that consumes up to 80% of the total cell energy(*2*), and its expression divergence (ED) is primarily due to the reduced gene expressions compared to the wild-type, it is likely that suppression of M5 *per se* saves energy, which promotes cell growth provided alterations of the other modules have already reduced the growth rate beneath a critical level. Consistent with these findings, deletion of *SSF1,* a member gene of M5, can be rescued by further deletion of *RPL16A,* a member gene of M3, or *PRM5,* a member gene of M6(*3*) (Fig. S9D). This finding challenges the common belief that down-regulation of ribosomal genes reduces the cell growth rate(*4–6*).

## Supplementary Note 2

A gene is said to affect a trait if deletion of the gene alters the trait. The common practice in current genetics considers only GIGs with statistically significant effects, but the idea of UGEs presumes that the statistically insignificant genetic effects of non-GIGs could be true signals. In fact, observation of the continuous distribution of the per-gene effect size suggests the limitation of using statistics to define genetic effects. We addressed this issue by analyzing the morphological information of individual cells of the yeast mutants. For each of the 4,718 mutants we compared the 501 traits between 50 mutant cells and 50 wild-type cells (Methods). We obtained a large number of both significant and insignificant genetic effects under the statistical cutoff of *p* < 0.001. As expected, with increasing mean effect size the frequency of significant effects increased substantially (Fig. S15A). The ubiquity hypothesis predicts that the difference between significant and insignificant effects may simply represent the variation of samplings from the same data population. To test this, we artificially modified the trait values of every wild-type cell by adding (or subtracting) a given effect size to form pseudo-mutants (Methods). The pseudo-mutant cells were then compared to the wild-type cells under the same statistical settings, and both significant and insignificant signals were observed for samplings from the same pseudo-mutant population that has true difference from the wild-type. Interestingly, the proportion of significant effects observed in the pseudo-mutants was similar to that of the real mutants (Fig. S15B), suggesting that the statistically insignificant signals of the yeast gene deletions can be well explained by true genetic effects. It is thus likely that every gene can show statistically significant impact on every trait provided with sufficiently large sample size and precise trait measurement.

---

**Table S1.**
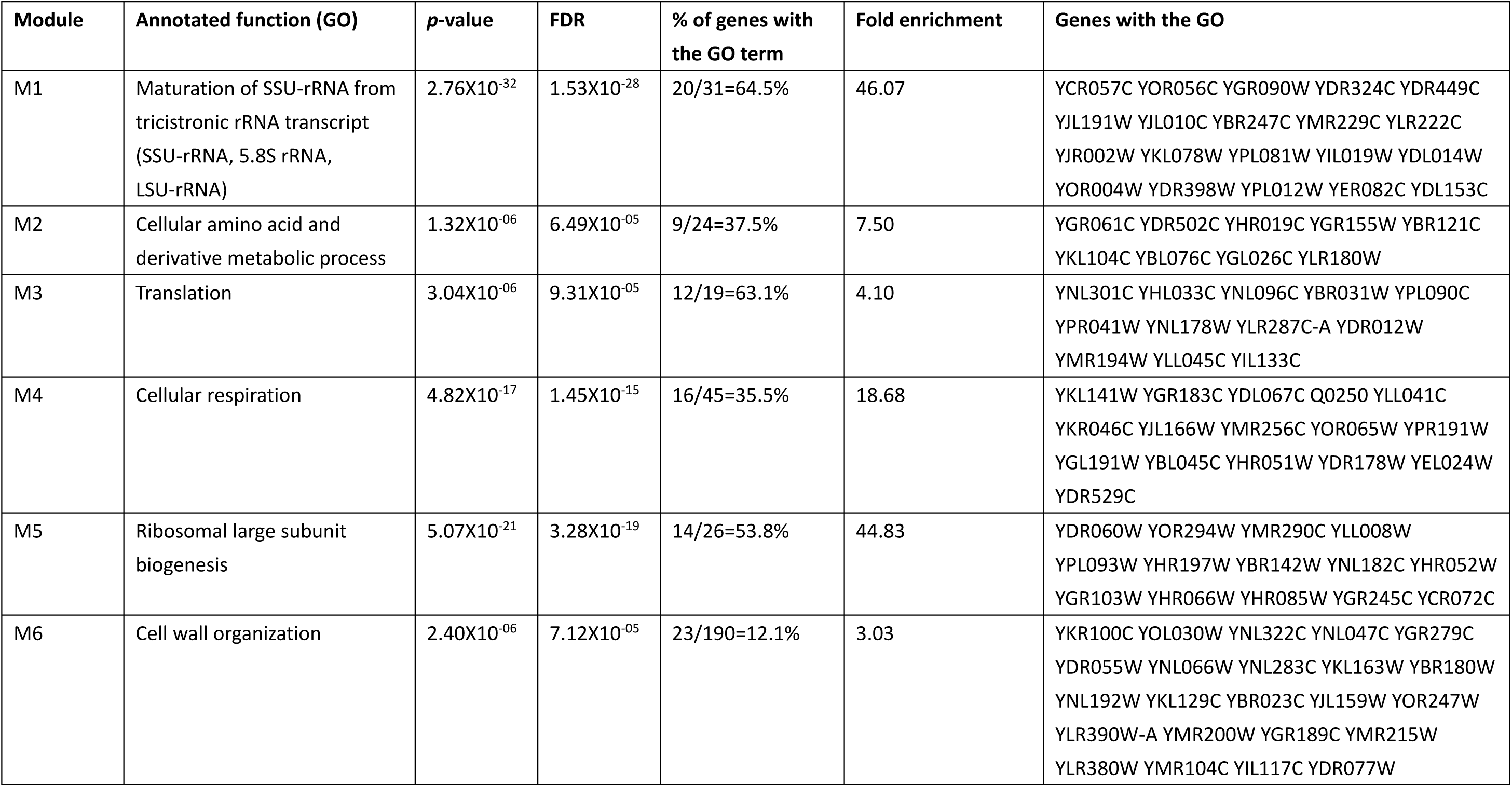
Characterization of the six growth-related EIG modules.

| Module | Annotated function (GO) | *p*-value | FDR | % of genes with the GO term | Fold enrichment | Genes with the GO |
| --- | --- | --- | --- | --- | --- | --- |
| M1 | Maturation of SSU-rRNA from tricistronic rRNA transcript (SSU-rRNA, 5.8S rRNA, LSU-rRNA) | 2.76X10<sup>-32</sup> | 1.53X10<sup>-28</sup> | 20/31=64.5% | 46.07 | YCR057C YOR056C YGR090W YDR324C YDR449C<br>YJL191W YJL010C YBR247C YMR229C YLR222C<br>YJR002W YKL078W YPL081W YIL019W YDL014W<br>YOR004W YDR398W YPL012W YER082C YDL153C |
| M2 | Cellular amino acid and derivative metabolic process | 1.32X10<sup>-06</sup> | 6.49X10<sup>-05</sup> | 9/24=37.5% | 7.50 | YGR061C YDR502C YHR019C YGR155W YBR121C<br>YKL104C YBL076C YGL026C YLR180W |
| M3 | Translation | 3.04X10<sup>-06</sup> | 9.31X10<sup>-05</sup> | 12/19=63.1% | 4.10 | YNL301C YHL033C YNL096C YBR031W YPL090C<br>YPR041W YNL178W YLR287C-A YDR012W<br>YMR194W YLL045C YIL133C |
| M4 | Cellular respiration | 4.82X10<sup>-17</sup> | 1.45X10<sup>-15</sup> | 16/45=35.5% | 18.68 | YKL141W YGR183C YDL067C Q0250 YLL041C<br>YKR046C YJL166W YMR256C YOR065W YPR191W<br>YGL191W YBL045C YHR051W YDR178W YEL024W<br>YDR529C |
| M5 | Ribosomal large subunit biogenesis | 5.07X10<sup>-21</sup> | 3.28X10<sup>-19</sup> | 14/26=53.8% | 44.83 | YDR060W YOR294W YMR290C YLL008W<br>YPL093W YHR197W YBR142W YNL182C YHR052W<br>YGR103W YHR066W YHR085W YGR245C YCR072C |
| M6 | Cell wall organization | 2.40X10<sup>-06</sup> | 7.12X10<sup>-05</sup> | 23/190=12.1% | 3.03 | YKR100C YOL030W YNL322C YNL047C YGR279C<br>YDR055W YNL066W YNL283C YKL163W YBR180W<br>YNL192W YKL129C YBR023C YJL159W YOR247W<br>YLR390W-A YMR200W YGR189C YMR215W<br>YLR380W YMR104C YIL117C YDR077W |

## Legends of supplementary figures

**Fig. S1.**
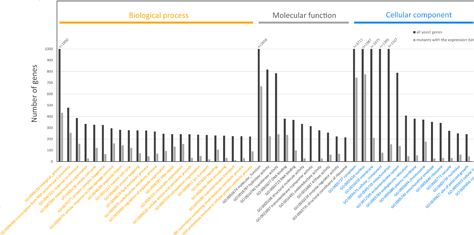
The >1,300 single-gene deletion mutants represent diverse genetic perturbations to the yeast cell.

**Fig. S2.**
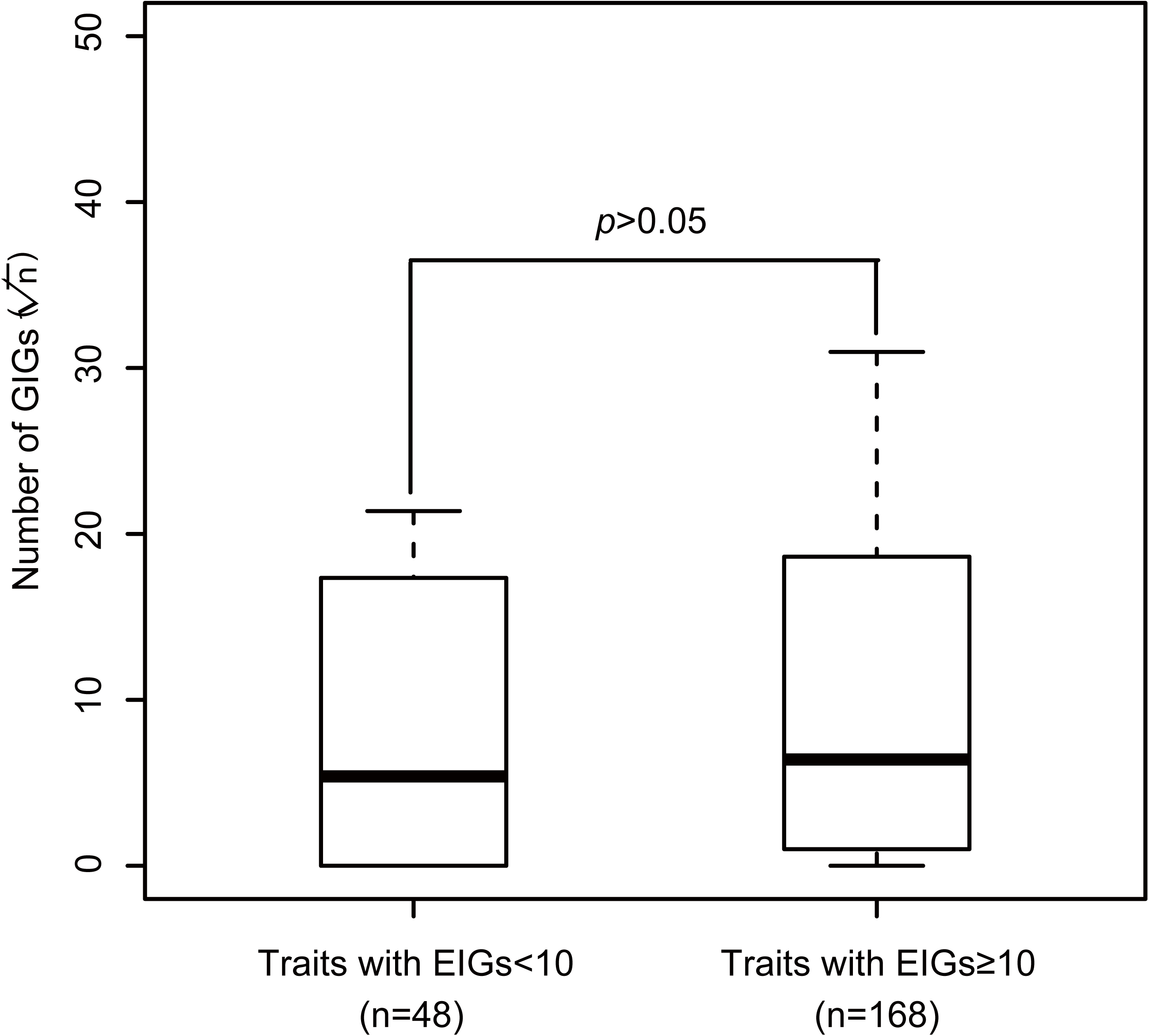
Traits with different numbers of EIGs have a comparable genetic complexity measured by their GIG numbers. Box-plots are presented, with the y-axis showing the square root of the number of GIGs. Mann-Whitney U test is used to compute the *p* value.

**Fig. S3.**
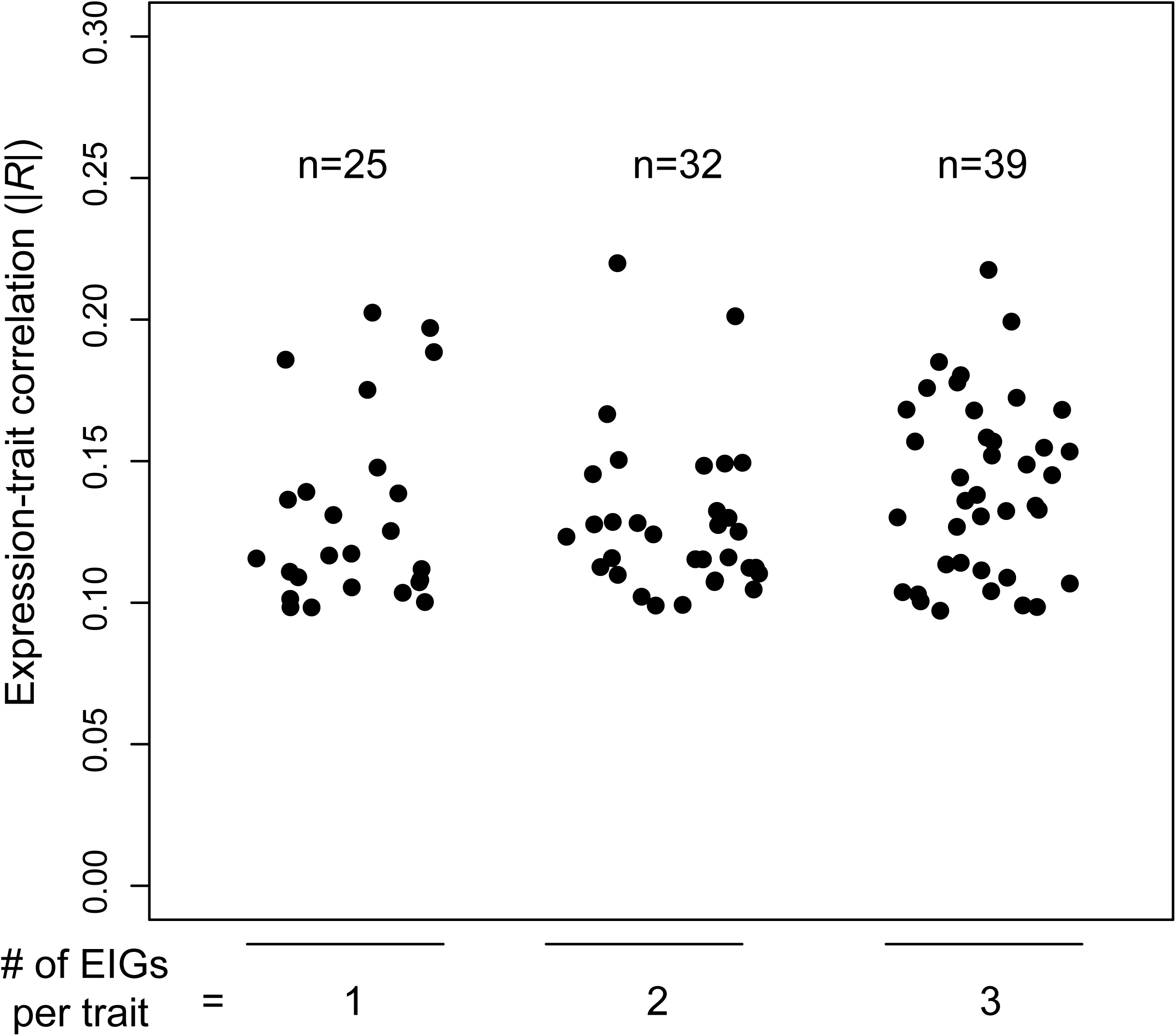
Out of the 118 EIG-trait correlations suitable for examination, 18 EIG ➔ trait and 27 trait ➔ EIG causal associations are reliably assigned. Thus, the proportion of causal EIGs ranges from 18/118 ~= 15% (assuming no positive in the 73 uncertain associations) to 18/(18+27) = 40%.

**Fig. S4.**
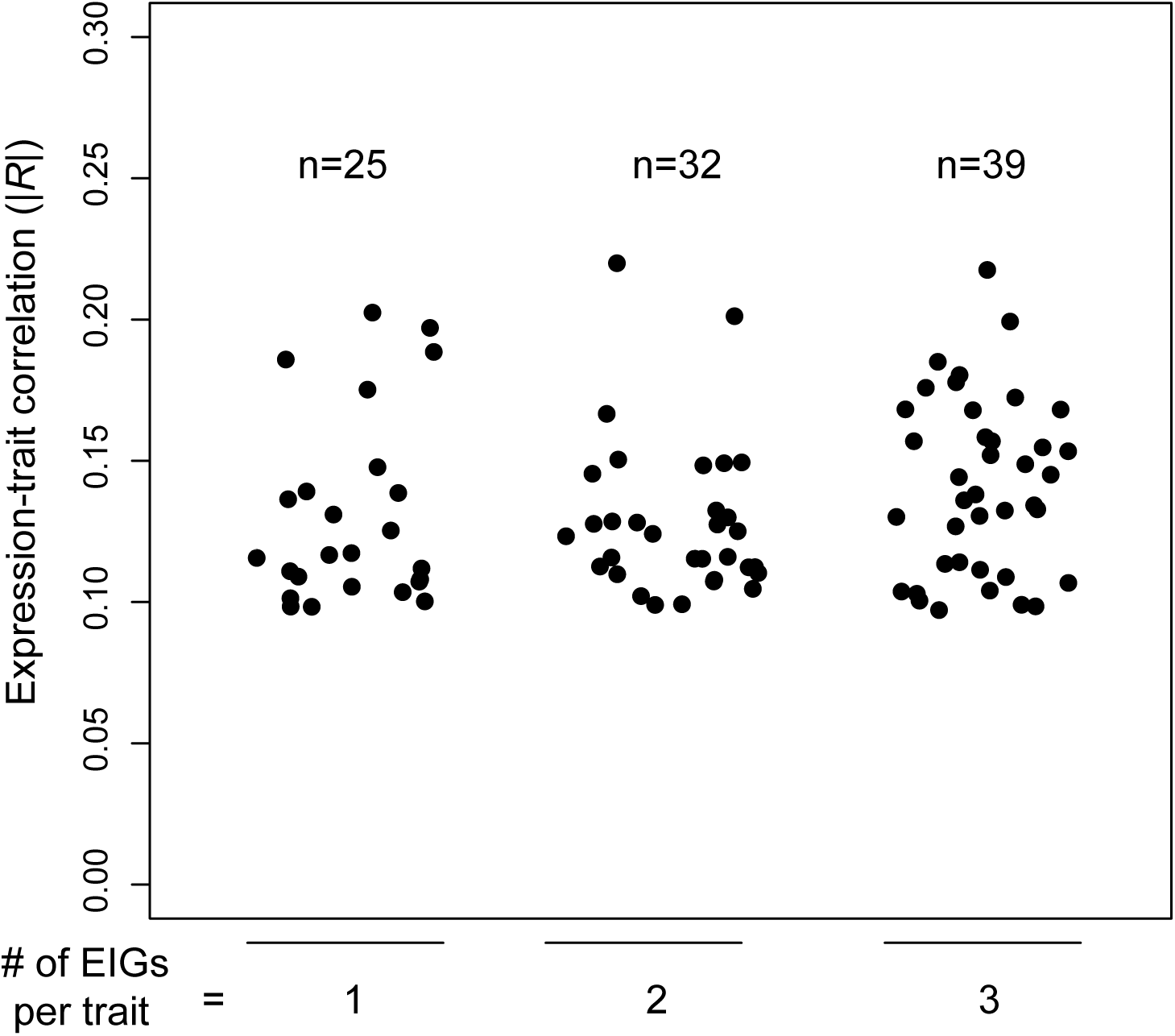
The variance explained by an individual EIG is minimal even in traits with only one, two, or three EIGs (x-axis). The y-axis shows the absolute value of the Pearson’s *R* between EIG expression and trait value in the Set #2 mutants. Each dot represents an EIG.

**Fig. S5.**
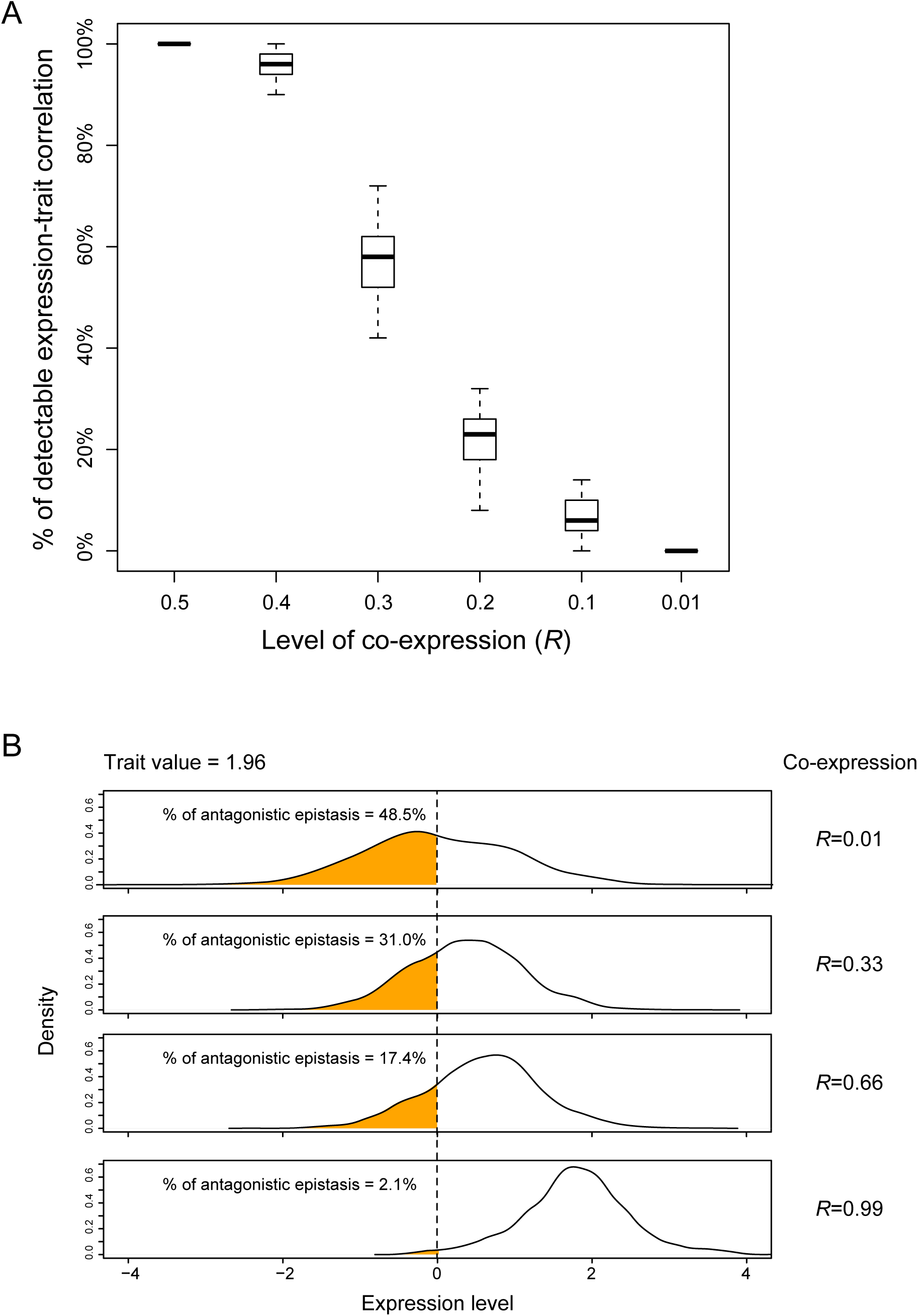
The coordination of causal factors determines the performance of observational approaches. **(A)** Reducing the co-expression of the 50 causal genes compromises the detection of a significant expression-trait correlation for every individual causal gene. The x-axis shows the average Pearson’s *R* of all gene pairs in 1,000 simulated expression profiles, and the y-axis is the proportion of causal genes that remain a significant expression-trait correlation (*q* < 0.01, n = 1,000, Pearson’s correlation analysis). **(B)** With a given trait value of 1.96, the proportion of down-regulation found for an individual causal gene is high when the co-expression is low. Because down-regulation of a causal gene alone should reduce the trait value to be negative (i.e., smaller than the wild-type), antagonistic epistasis has to be invoked to explain such down-regulations when the trait value is positive (1.96).

**Fig. S6.**
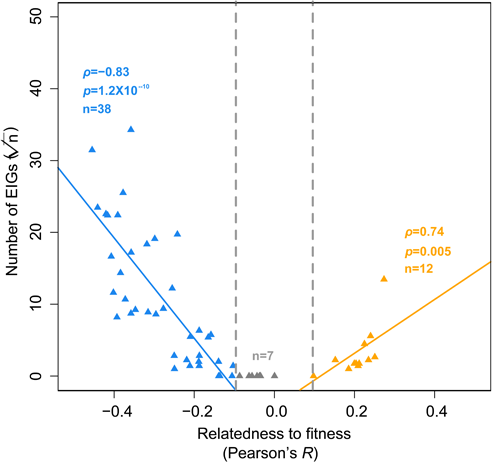
Same as Fig. 1, except that the 57 exemplar traits are considered.

**Fig. S7.**
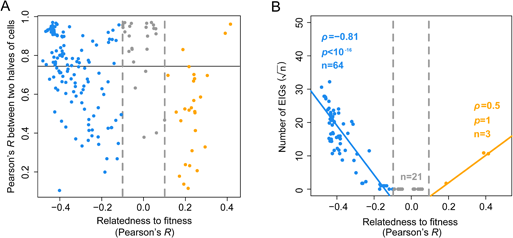
The varied quality of trait measures cannot explain the reduced number of EIGs in traits less coupled with fitness. **(A)** The y-axis shows the Pearson’s *R* of the trait values between the two halves of cells examined for each mutant, and the horizontal line marks *R* = 0.75. Each dot represents a trait, and a total of 216 traits with the information of individual cells are included. **(B)** Same as Fig. 1, except that the 88 traits with good internal consistency (*R* > 0.75) between the two halves of examined cells are considered.

**Fig. S8.**
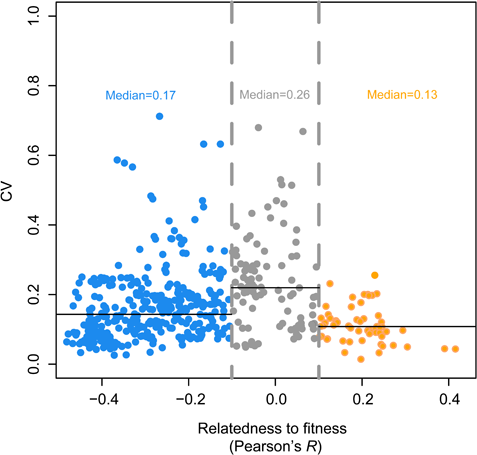
The between-mutant CV of traits with no significant correlation to fitness is not smaller than that of fitness-coupled traits, where CV stands for coefficient of variation, suggesting that the reduced number of EIGs in traits less coupled with fitness cannot be explained by the lack of variation.

**Fig. S9.**
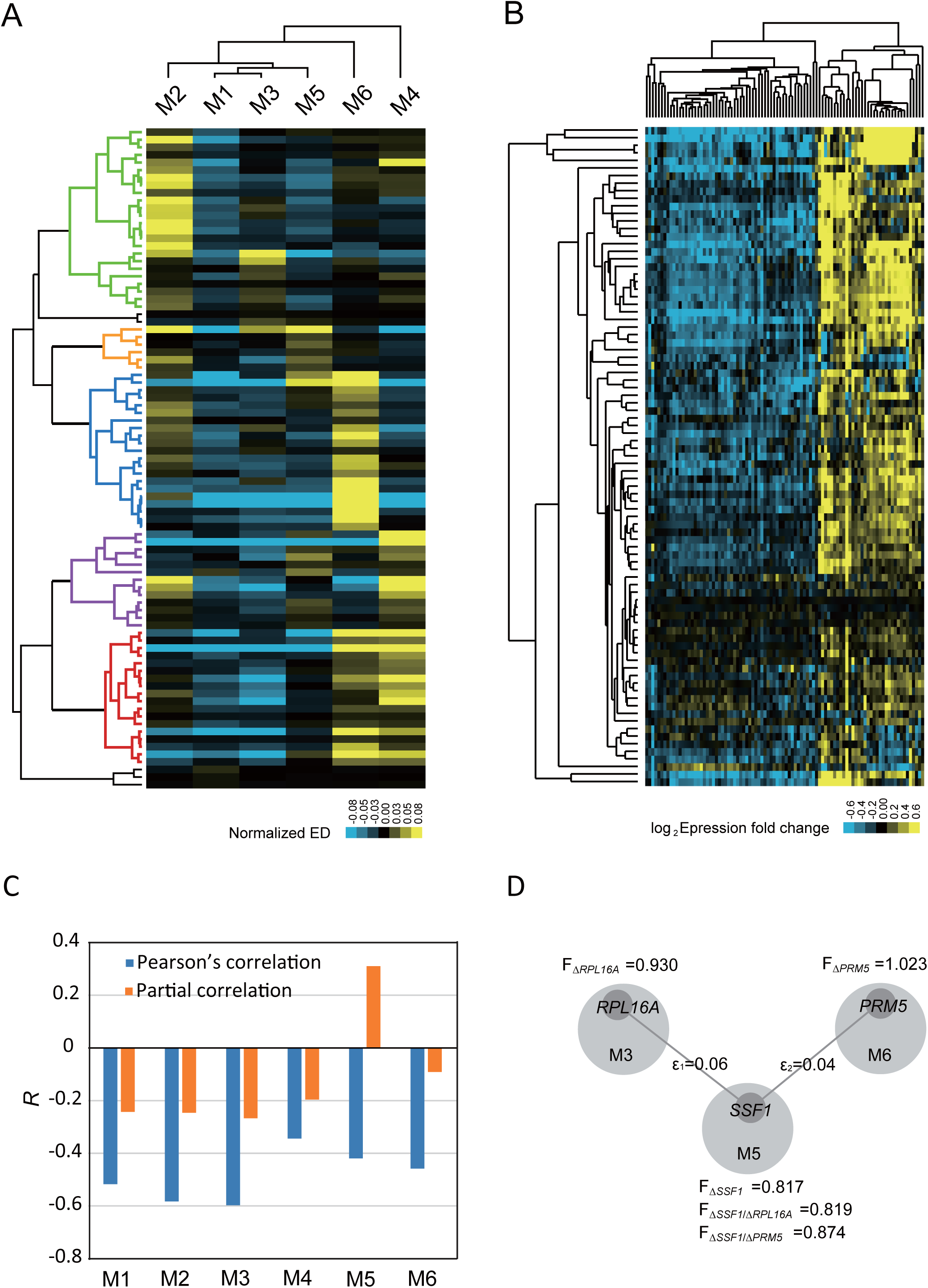
Novel mechanistic insights on yeast cell growth provided by the six EIG-modules. **(A)** The five types of growth defects defined by the six EIG-modules. Each row represents a slow-growth mutant, and the expression distance (ED) of a module is normalized by subtracting its mean ED in the 87 mutants. **(B)** No clear mutant cluster is found based on expressions of individual genes of the six modules. Each row represents a mutant and each column represents a gene, with the expression changes relative to the wild-type being shown. **(C)** The Pearson’s *R* between module activity and cell growth rate for each of the six EIG-modules, in comparison to that of the partial correlation that controls for the other five modules. **(D)** The rescuing epistasis between *SSF1* of M5 and *PRL16A* of M3 or *PRM5* of M6. F represents the relative growth rate (or fitness) of a mutant, with ε_1_ = F_Δ_*_SSFI/_*_Δ_*_PRL16A_* − F_Δ_*_SSF1_* × F_Δ_*_PRL16A_* and ε_2_ = F_Δ_*_SSF1/_*_Δ_*_PRM5_* − F_Δ_*_SSF1_* × F_Δ_*_PRM5_*.

**Fig. S10.**
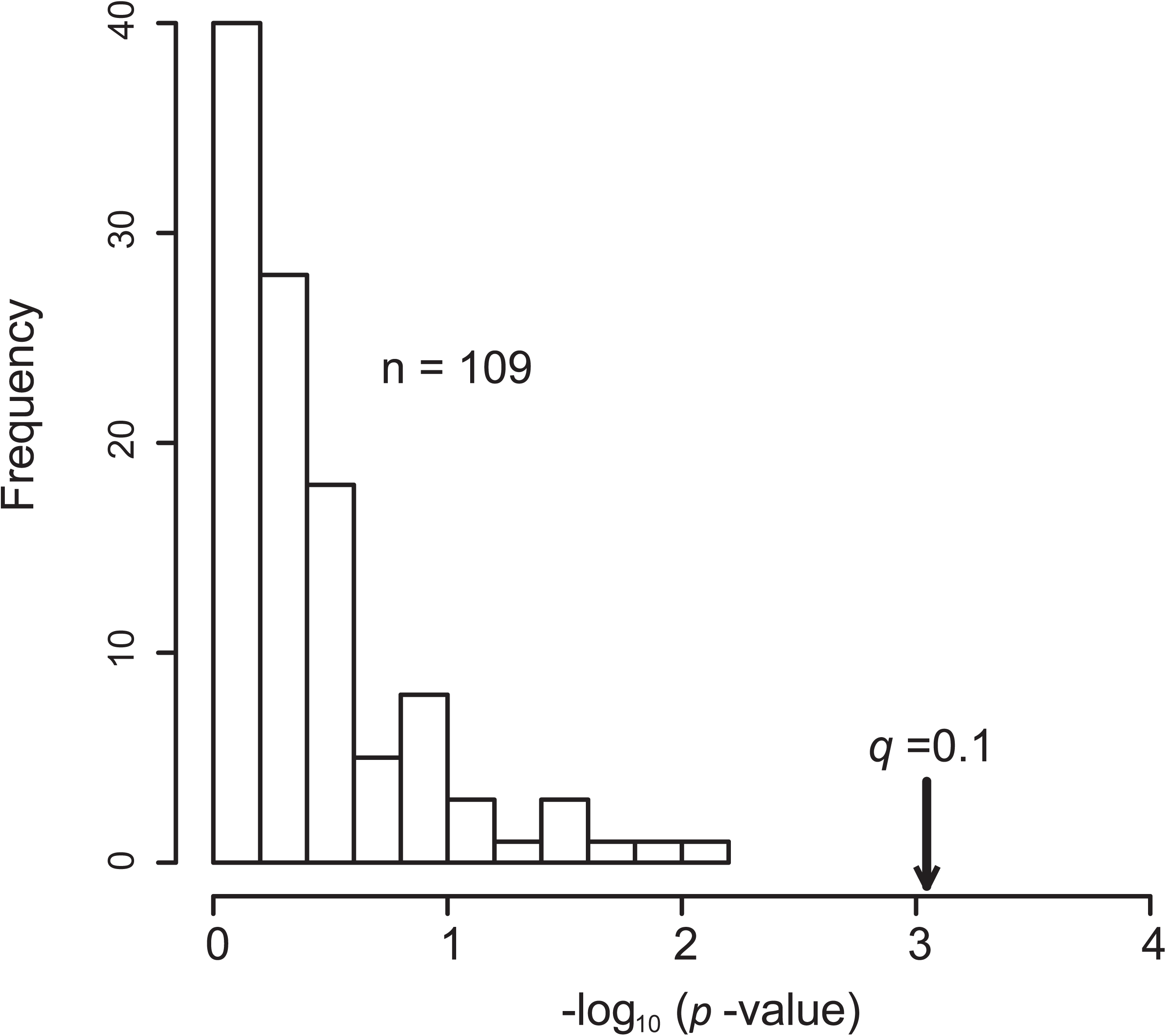
No significant overlaps between EIGs and GIGs of the same traits in a total of 109 traits each with at least 10 EIGs and 10 GIGs. Chi-square test is used to compute the *p*-values shown at the x-axis, and *q*=0.1 shows the expected significance cutoff after controlling for multiple testing.

**Fig. S11.**
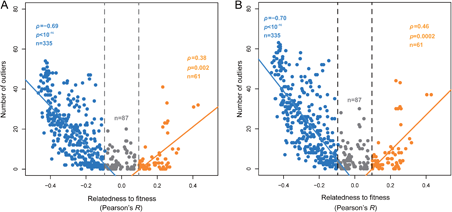
Same as Fig. 4E, except that the Z-score cutoff for identifying outliers is reduced to 4.60 **(A)** and 4.26 **(B**), corresponding to *q* smaller than 0.005 and 0.01, respectively.

**Fig. S12.**
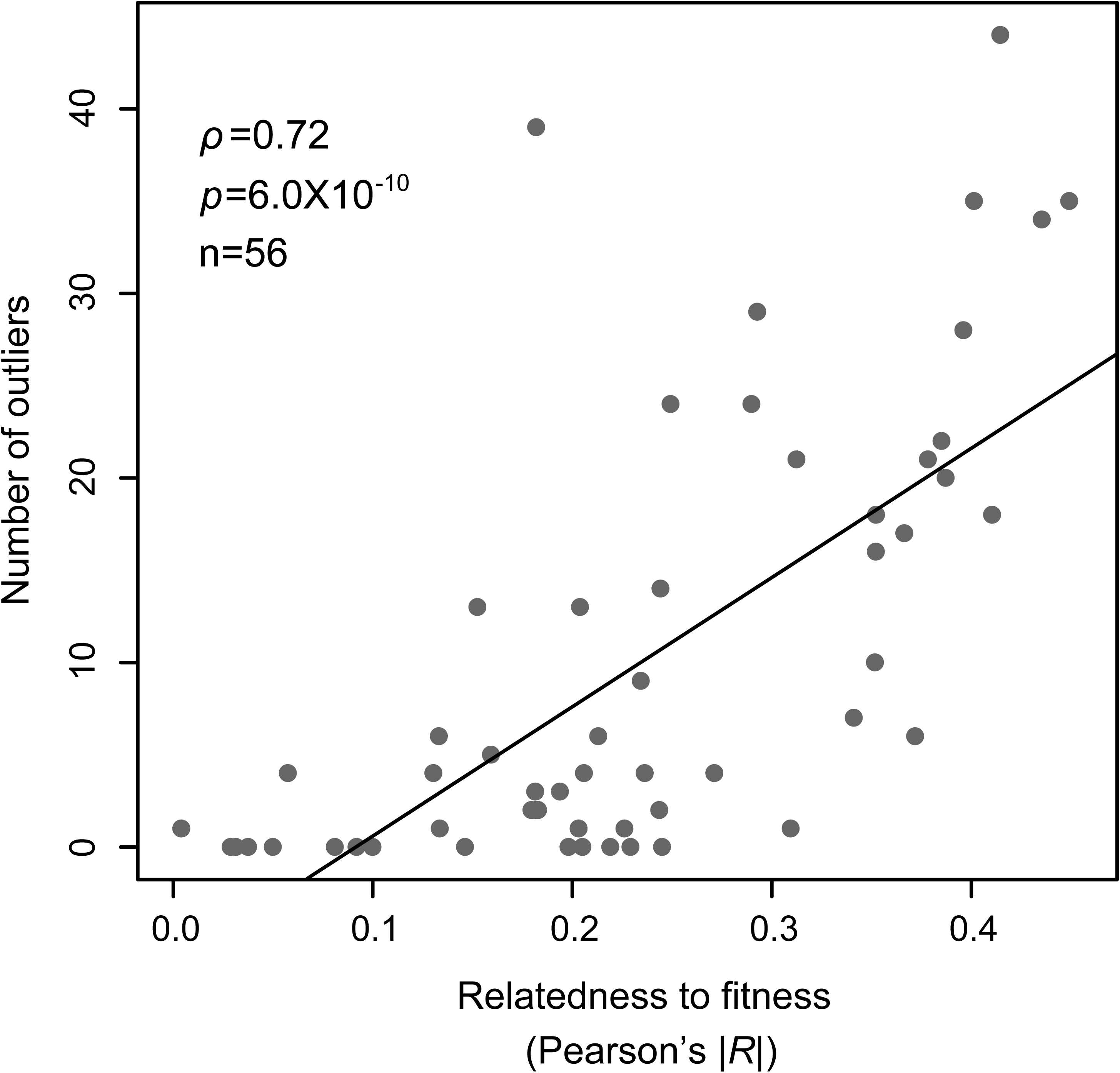
Same as Fig. 4E, except that only the 56 exemplar traits are considered, and that the absolute trait relatedness to fitness is shown at the x-axis.

**Fig. S13.**
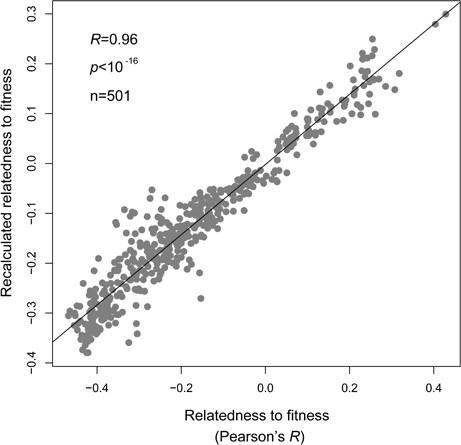
The recalculated trait relatedness to fitness (y-axis) after removing the effects of outliers is highly correlated to the original one (x-axis), indicating that fitness coupling is the cause of its correlation with the number of outliers.

**Fig. S14.**
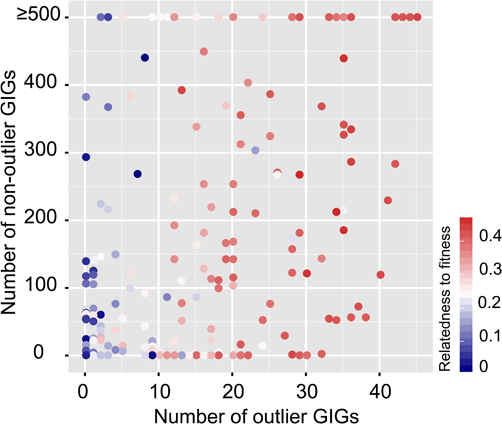
The number of outlier GIGs and non-outlier GIGs as a function of the trait relatedness to fitness. Each dot represents a trait, and a total of 216 traits are included.

**Fig. S15.**
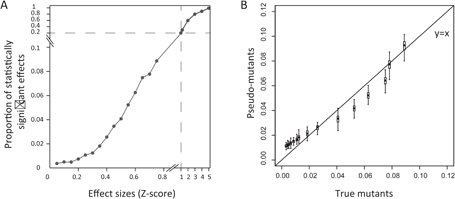
Both statistically significant and insignificant signals can be explained by true genetic effects. **(A)** The larger mean effect size a mutant has, the higher probability that significant differences between 50 mutant cells and 50 wild-type cells are observed. **(B)** The probability of observing significant signals in true mutants is similar to that in the simulated pseudo-mutants. We considered only effect sizes ranging from Z = 0 to Z = 0.77, which covers 50% of the data with *Z* > 0 in a standard Gaussian distribution. The box and error bar encompass 50% and 90% of the data derived from 100 simulations, respectively.

**Fig. S16.**
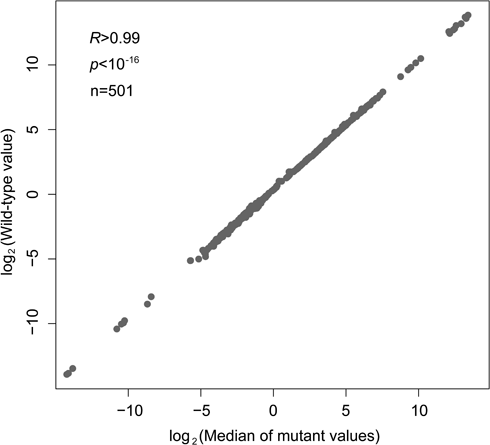
The distribution of trait values of the 4,718 mutants is bell-shaped, with the median nearly equivalent to the trait value of the wild-type for nearly all of the 501 morphological traits.

**Fig. S17.**
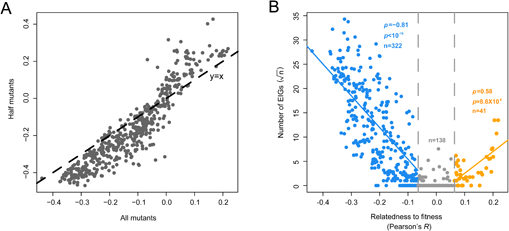
**(A)** The trait relatedness to fitness estimated from half of the mutants is highly correlated to that estimated from all mutants. **(B)** Same as Fig. 1, except that the trait relatedness to fitness is estimated from all mutants.

